# H_2_-driven xylitol production in *Cupriavidus necator* H16

**DOI:** 10.1101/2024.10.22.619622

**Authors:** Tytti Jämsä, Nico J. Claassens, Laura Salusjärvi, Antti Nyyssölä

## Abstract

**Background:** Biocatalysis offers a potentially greener alternative to chemical processes. For biocatalytic systems requiring cofactor recycling, hydrogen emerges as an attractive reducing agent. Hydrogen is attractive because all the electrons can be fully transferred to the product, and it can be efficiently produced from water using renewable electricity. In this article, resting cells of *Cupriavidus necator* H16 harboring a NAD-dependent hydrogenase were employed for cofactor recycling to reduce D-xylose to xylitol, a commonly used sweetener. To enable this bioconversion, D-xylose reductase from *Scheffersomyces stipitis* was heterologously expressed in *C. necator*.

**Results:** D-xylose reductase was successfully expressed in *C. necator*, enabling almost complete bioconversion of 30 g/L of D-xylose into xylitol. It was found that over 90% of the energy and protons derived from hydrogen were spent for the bioconversion, demonstrating the efficiency of the system. The highest xylitol productivity reached was 0.7 g L^-1^ h^-1^. Additionally, the same chassis efficiently produced L-arabitol and D-ribitol from L-arabinose and D-ribose, respectively.

**Conclusions:** This study highlights the efficient utilization of renewable hydrogen as a reducing agent to power cofactor recycling. Hydrogen-oxidizing bacteria, such as *C. necator*, can be promising hosts for performing hydrogen-driven biocatalysis.

## Background

Biocatalysis is increasingly applied across different industries due to its efficiency and environmental benefits compared to chemical transformations. Biocatalysis using oxidoreductases often requires cofactors, such as nicotinamide nucleotides NADH and NADPH, but their stoichiometric addition to reaction mixtures is not economically feasible (1). Therefore, cofactor recycling systems are required. Traditionally cofactor recycling of NAD(P)H is performed using sacrificial substrates, such as glucose and formate, which are oxidized during the biocatalysis, while the cofactor is regenerated to its reduced form (2). This is not carbon-efficient, since D-gluconolactone and carbon dioxide (CO_2_) are formed as by-products from glucose and formate, respectively. Specifically for the by-product D-gluconolactone, a substantial number of energy-rich electrons are wasted. In addition, glucose is produced by agriculture, which can decrease the overall sustainability of the production process due to, for example, competition with food production and environmental burdens of agriculture. Formate can be made from CO_2_ and renewable electricity using electrochemical reduction. However, electrochemical production of formate has not yet been scaled-up and is not as energy-efficient as the electrochemical production of hydrogen (H_2_) from water and electricity, which is already performed on a large scale (3).

H_2_ is an attractive, byproduct-free sacrificial substrate for cofactor recycling. Molecular H_2_ can be oxidized by organisms using hydrogenases. Some of the most extensively researched H_2_-uptake hydrogenases are found in the hydrogen-oxidizing bacterium *Cupriavidus necator* H16 (formerly *Ralstonia eutropha*) (4). This bacterium possesses two types of oxygen-tolerant hydrogenases that provide the cells with reducing power: membrane-bound and soluble hydrogenases (5). The membrane-bound hydrogenase is located in the cytoplasmic membrane where it directly feeds electrons to the respiratory chain for ATP production by oxidative phosphorylation. The NAD-dependent hydrogenase resides in the cytoplasm and is therefore referred to as soluble hydrogenase (SH) (6, 7). The electrons and protons from H_2_ can be directly transferred to NAD^+^ by the SH, reducing NAD^+^ to NADH while simultaneously oxidizing hydrogen (4).

*C. necator* H16 SH has been researched broadly for cofactor recycling using purified enzymes (8-12). Compared to purified enzymes, whole-cell biocatalysts can provide enzyme stabilization, as isolated intracellular enzymes typically lose activity more quickly than when they are in their natural cellular environments (13). Whole-cell systems can also have lower production costs by eliminating the need for enzyme purification or external addition of cofactors. Additionally, whole cells are generally more robust and exhibit better tolerance to inhibitors compared to purified enzymes. Whole cells were used in one of the earliest studies of SH-catalyzed cofactor recycling in the 1980s where native *C. necator* cells reduced CO_2_ to formate with a 30% yield (14). More recently, whole cells of recombinant *C. necator* have been employed as a biocatalyst by Oda et al. (15) and Assil-Companioni et al. (16) for reduction of hydroxyacetone to (*R*)-1,2-propanediol and for asymmetric C=C bond reduction of unsaturated cyclic ketones, respectively. However, given the limited extent of research and number of products, further studies are required to investigate *C. necator* as a whole-cell biocatalyst. In the current study, we show that the scope of hydrogen-driven reductive bioconversions can be broadened to encompass a new class of products, sugar alcohols, with xylitol as a particular example.

Xylitol (C_5_H_12_O_5_) is a sugar alcohol used widely as a sweetener (17). It is currently produced chemically from D-xylose (C_5_H_10_O_5_), which is the second most abundant sugar in lignocellulosic biomass (18). The chemical production of xylitol requires extensively purified D-xylose to avoid inactivation of the catalyst. In contrast, biotechnological production does not have this requirement and exhibits greater tolerance for inhibitors. Many yeasts, along with some bacteria and filamentous fungi, naturally reduce D-xylose to xylitol (18). However, numerous studies have focused on heterologous expression of D-xylose reductase (XR), aiming to increase yield and productivity of xylitol (19-23). All these studies use sugars to power the conversion, and to our knowledge H_2_ has not been used as the electron and energy source for the D-xylose-to-xylitol conversion.

The objective of this study was to develop a whole-cell biocatalyst strain of *C. necator* for xylitol production using H_2_ as the electron donor. To achieve this, XR from *Scheffersomyces stipitis* (formerly *Pichia stipitis*) was expressed in *C. necator* to allow the cells to convert D-xylose into xylitol (Figure 1). The native SH of *C. necator* enabled the H_2_-driven cofactor recycling required for bioconversion. Resting (i.e., non-dividing) cells were used in the experiments. These viable cells exhibit reduced metabolic activity, which allows energy to be directed towards bioconversion instead of biomass accumulation (24). With the experimental set-up used, we achieved nearly full quantitative conversions.

**Figure 1.**
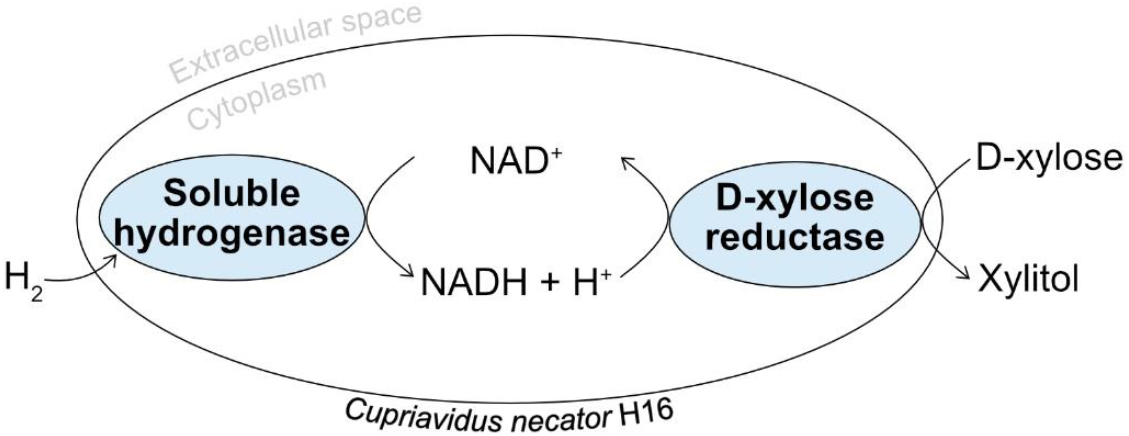
H_2_-driven production of xylitol from D-xylose in whole cells of *Cupriavidus necator* H16 expressing D-xylose reductase from *Scheffersomyces stipitis*.

## Materials and methods

### Strains and culture media

Bacterial strains and plasmids used in this work are listed in Table 1 and the primers in Table S1 (see Additional file 1). *C. necator* H16 strains were grown in either rich BD BBL™ Trypticase™ Soy Broth (TSB) or minimal AUT medium (25) supplemented with 2 g/L fructose and glycerol (FG). Compared to the original AUT medium, the amount of NiCl_2_ was doubled and SL-6 trace elements were added (1:1000) (26). *Escherichia coli* DH10β was used for plasmid construction. Tetracycline was added at a concentration of 10 μg/mL for *E. coli* and 5 μg/mL for *C. necator* when required. The sugars and sugar alcohols used in the study were purchased from Sigma-Aldrich.

**Table 1.** Bacterial strains and plasmids used in the study. The strains with △*phaC* deletion are unable to produce polyhydroxybutyrate (PHB) for carbon and energy storage.

| Strain | Relevant characteristics | Source or reference |
| --- | --- | --- |
| *C. necator* H16 $\Delta phaCAB$ | $\Delta phaC1 \Delta phaA1 \Delta phaB1$ | (27) |
| *C. necator* H16 $\Delta phaC1$ | $\Delta phaC1$ | (28) |
| *C. necator* H16 $\Delta A0006$ | $\Delta phaC1 \Delta A0006$ | This study |
| *C. necator* H16 $\Delta phaCAB\_gfp$ | $\Delta phaCAB$ derivative, pPj5:GFP | This study |
| *C. necator* H16 $\Delta phaC\_gfp$ | $\Delta A0006$ derivative, pPj5:GFP | This study |
| *C. necator* H16 $\Delta phaCAB\_xr$ | $\Delta phaCAB$ derivative, pPj5:XRsti | This study |
| *C. necator* H16 $\Delta phaC\_xr$ | $\Delta A0006$ derivative, pPj5:XRsti | This study |
| **Plasmid** |  |  |
| pLO3 | Suicide vector, sacB | (29) |
| pLO3-*A0006* | pLO3 derivative with 1100 bp upstream and downstream regions of *A0006* | This study |
| pPj5:GFP | pSEVA521 containing Pj5 promoter and GFP | (30) |
| pPj5:XRsti | pPj5:GFP derivative, GFP replaced with D-xylose reductase from *Scheffersomyces stipitis* (XRsti) | This study |

### Strain construction

XR from *Scheffersomyces* (*Pichia*) *stipitis* was ordered codon-optimized for *C. necator* from GenScript (Additional file 1: Table S2) and PCR amplified with Q5 High-Fidelity 2X Master Mix (NEB). The backbone plasmid pSEVA521+Pj5:GFP and the amplified insert were digested with *Spe*I-HF and *Hin*dIII-HF (NEB) and ligated using T4 DNA ligase (NEB) to gain pPj5:XRsti. The verified plasmid was transformed into electrocompetent *C. necator* cells. For preparing competent cells, *C. necator* was grown in 100 mL of TSB supplemented with 20 mM of fructose to an optical density at 600 nm (OD_600_) of 0.6 and washed twice with 1 mM MgSO_4_. The pellet was resuspended into 2 mL of 1 mM MgSO_4_ and 1 mL of 60% glycerol. Aliquots (50 μL) were stored in -80°C. Cells were mixed with plasmids (250 ng) in a 0.2 cm electroporation cuvette (Bio-Rad), incubated 10 min on ice and electroporated with Electro Cell Manipulator ECM®630 (BTX) with the following settings: 2.5 kV, 200 Ω and 25 μF. Super Optimal Broth with 20 mM fructose (950 μL) was added immediately after electroporation and cells were incubated at 30°C and 180 rpm for 2-3 hours before plating on BBL™ Trypticase™ Soy Agar (TSA).

*C. necator* H16 △*A0006* was constructed from *C. necator* H16 △*phaC1* by deleting *A0006* with pLO3-based suicide vector as previously described (29) (Additional file 1: Table S3). The △*A0006* restriction enzyme knockout increases electroporation efficiency (31, 32) and is not expected to have any metabolic effects.

### XR activity assay

For the XR activity assay, cells were grown overnight in TSB and then harvested by centrifugation (10 min, 2800 g). The pellet was washed once with 50 mM potassium phosphate buffer (pH 7.5), resuspended into 1 mL of the same buffer supplemented with cOmplete™ protease inhibitor (Roche) and moved into a 2 mL screw-cap tube with 400 μL of 0.5 mm diameter glass beads. The cells were disrupted with FASTPREP-24 5G (MP Biomedicals) for 2×30sec at 6 m/s speed. After disruption, the tube was centrifuged for 10 min at 16000 g, and the supernatant (soluble extract) was collected. Total protein concentration was analyzed from the soluble extract by Quick Start™ Bradford Protein Assay (Bio-Rad) using bovine serum albumin as the standard. XR activity assays were conducted in 96-well plates. The reaction mixture (330 μL) contained 0.15 mM of cofactor (NADH or NADPH), 50 mM potassium phosphate buffer (pH 6.0), 200 mM D-xylose and an appropriate amount of soluble extract. Absorbance was measured at 340 mM using Epoch 2 Microplate Spectrophotometer (BioTek). One unit of xylose reductase activity was defined as μmol of NAD(P)H oxidized per minute. Specific activities were expressed as units per milligram of total protein. The results are given as averages of triplicate assays.

### Bioconversions with resting cells

Precultures were grown overnight in TSB media. FG media with tetracycline was inoculated to an initial OD_600_ of 0.1 and grown for three days, reaching a final OD_600_ of approximately 4. The culture was centrifuged for 10 min at 2800 g and washed twice with 100 mM sodium phosphate buffer (pH 7.0) to remove carbon and nitrogen sources. The cells were resuspended in the same buffer with 30 g/L of substrate to an OD_600_ of 13-17, if not mentioned otherwise. This range of OD_600_ corresponds approximately to a cell dry weight of 4.4-5.4 g/L. The prepared cell suspension (5 mL) was transferred into an anaerobic serum bottle with a rubber stopper. D-xylose was the primary substrate, but one bioconversion was also performed using L-arabinose and another using D-ribose. Three replicate bottles per bioconversion condition were prepared.

For the first bioconversion experiments, 100 mL serum bottles were filled with H_2_ using vacuum-gas cycles to reach specific H_2_ and oxygen (O_2_) concentrations. As negative controls, bioconversions were performed under ambient air. The bottles were incubated at 30°C and 150 rpm. Samples (200 μL) were taken by opening the rubber stopper and the bottles were refilled with gasses after sampling. For bioconversion optimization, 50 mL serum bottles were filled with H_2_ at the start of the bioconversion by flushing with 100% H_2_ at 0.5 mL min^-1^ for 2 minutes. Samples (200 μL) were taken using a needle and syringe through the rubber stopper of sealed bottles. The approximate H_2_ gas consumption was measured after bioconversions by filling a 50 mL syringe with air and recording the volume of air aspirated into the bottle through the syringe needle. Samples were centrifuged for 10 min at 16000 g and the supernatants were analyzed for sugars and sugar alcohols.

### Calculation of H_2_ consumption

The amount of H_2_ consumed was calculated from the approximated H_2_ gas consumption (Equation 1):

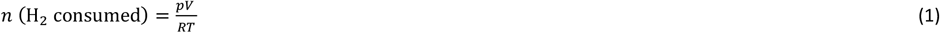

where *p* is the pressure (1 bar), *V* is the gas volume consumed (L), *R* is the gas constant (0.08314 L bar K^-1^ mol^-1^) and *T* is the temperature (298.15 K).

One mole of H_2_ is needed to reduce one mole of xylose (Equation 2):

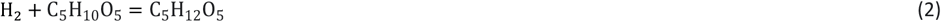

The total amount of xylitol produced was calculated as the sum of the xylitol at the end of the bioconversion and the xylitol in the fractions taken out during sampling. The ratio of xylitol production to H_2_ consumption was calculated to determine the proportion of energy and protons transferred from H_2_ to xylitol.

### Analysis of sugars and sugar alcohols

Two different high performance liquid chromatography (HPLC) systems were used for D-xylose and xylitol analysis: Prominence-i LC-2030C (Shimadzu) equipped with Hi-Plex H 7.7×300 mm column (Agilent) at 45°C and 10 mM H_2_SO_4_ as eluent at a flow rate of 1 ml min^-1^ and Vanquish Flex (Thermo Fisher Scientific) equipped with Aminex Fast Acid Analysis and HPX-87H columns (Bio-Rad) at 55°C and 2 mM H_2_SO_4_ as the eluent at a flow rate of 0.5 ml min^-1^. An injection volume of 10 μL was used in both HPLCs and the compounds were detected with a refractive index detector. L-arabinose, L-arabitol, D-ribose and ribitol were analyzed with high pressure ion chromatography (HPIC) Dionex ICS-6000 (Thermo Fisher Scientific) with CarboPac PA20 column (Thermo Fisher Scientific) at 30°C and 10 mM KOH as the eluent at a flow rate of 0.5 ml min^-1^. An injection volume of 2.5 μL was used and the compounds were detected with an electrochemical detector. Conversion yields were calculated by dividing the final molar concentration of xylitol by the initial molar concentration of xylose.

## Results

### Xylose reductase from *S. stipitis* is functionally produced in *C. necator* H16

In this study, two *C. necator* strains, △*phaC* and △*phaCAB*, with partial or full knockouts of the native pathway for storage polymer polyhydroxybutyrate (PHB) formation, were used to avoid the accumulation of this by-product. Before constructing the XR expressing strains, it was confirmed that the host strains cannot grow on C5 sugars and sugar alcohols used in the study (Additional file 1: Fig. S1). Both strains were then transformed with the plasmids pPj5:XR, carrying the codon-optimized XR from *S. stipitis*, and pPj5:GFP, as a negative control. Strains were cultivated heterotrophically, and their soluble extracts were tested for XR activity. The soluble extracts of both XR strains showed reductase activity with both NADPH and NADH cofactors, whereas no activity was detected in the negative control strains (Table 2). Codon-optimized XR from *Candida parapsilosis* was also expressed in the host strains, but no activity was detected (data not shown).

**Table 2.** Specific D-xylose reductase activities of soluble extracts with NADPH or NADH as cofactors (U/mg). The average from three measurements is shown with the standard deviation. No activity could be detected in the negative controls.

| Strain | NADPH | NADH |
| --- | --- | --- |
| $\Delta phaCAB\_xr$ | 0.71±0.20 | 0.48±0.01 |
| $\Delta phaC\_xr$ | 0.49±0.01 | 0.41±0.03 |

### Xylose is fully converted to xylitol by *C. necator* △*phaCAB*

The first bioconversion experiment was performed at 30 g/L D-xylose by both *C. necator* strains: △*phaCAB_*xr and △*phaC_*xr. The suspensions were incubated under H_2_ (85% H_2_ + 15% air) or 100% ambient air. Under H_2_, the △*phaCAB_*xr strain reached almost complete bioconversion to xylitol within 16 days, whereas the △*phaC_*xr strain converted 74% of the provided D-xylose at the same time (Figure 2). Therefore, further bioconversions were performed using the △*phaCAB_*xr strain. D-xylose was also converted to xylitol in the absence of H_2_ by both strains, but the conversion yields were under 21% after 16 days. This demonstrated successful cofactor recycling in resting cells of *C. necator* using H_2_.

**Figure 2.**
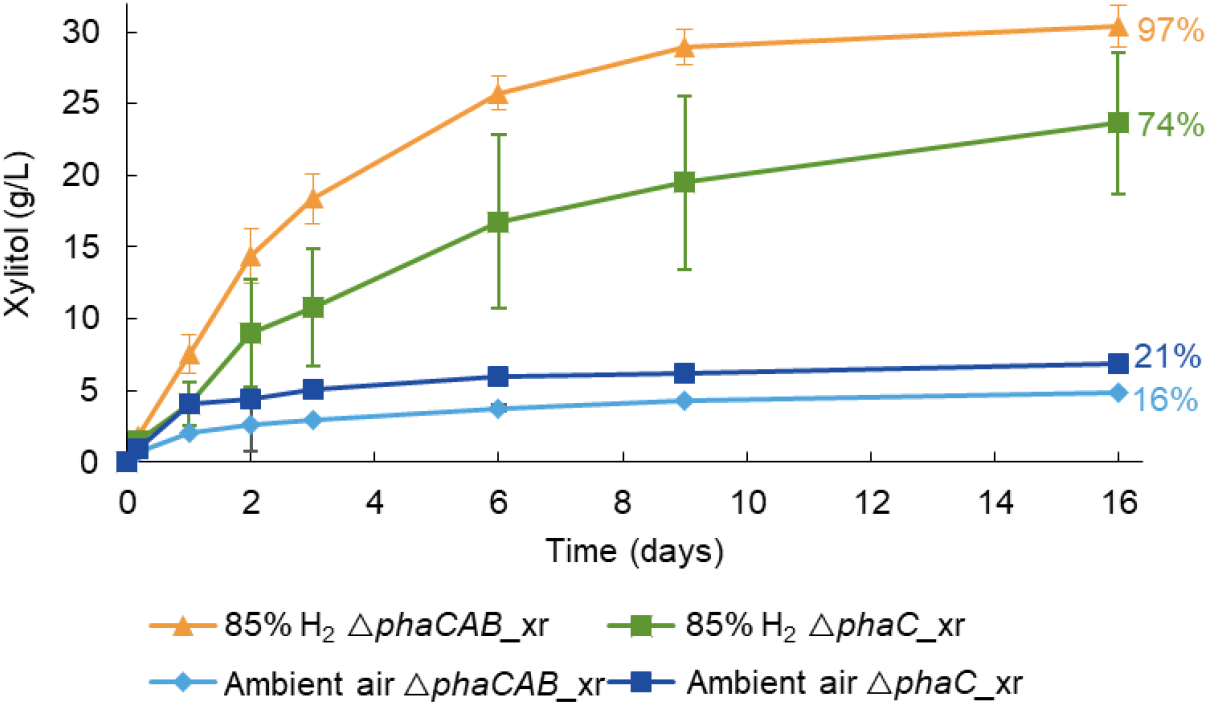
H_2_-driven production of xylitol by *C. necator* △*phaCAB_*xr and △*phaC_*xr. The initial concentration of D-xylose was ∼30 g/L in 100 mM sodium phosphate buffer (pH 7.0) with the cells at OD_600_ of 17 (△*phaCAB_*xr) and 15 (△*phaC_*xr). The used gas mixture consisted of 85% H_2_, 12% N_2_, and 3% O_2_ while ambient air contains 78% N_2_ and 21% O_2_. The average from three bioconversions is shown with the standard deviation. The percentages represent the final conversion yields, calculated by dividing the xylitol concentration measured at the final time point by the xylose concentration at the start of the experiment.

### Oxygen is not required for the bioconversion

Since oxygen, acting as an electron acceptor, can provide the cells with energy via oxidative phosphorylation, we examined the effect of oxygen concentration on the conversion rate. Three different oxygen concentrations (0, 1, and 4%) were tested. The conversion rates and yields showed little variation between the different oxygen concentrations (Figure 3), with over 85% conversion yields being reached within 10 days in all conditions. After confirming that oxygen was not required for the bioconversion, it was tested whether the amount of H_2_ was a limiting factor. Bioconversions with multiple H_2_ flushes during the experiment were compared to bioconversions with a single H_2_ flush at the start. The results showed that multiple H_2_ flushes failed to improve the conversion yield (Additional file 1: Fig. S2). Therefore, further bioconversions were conducted with only an initial H_2_ flush.

**Figure 3.**
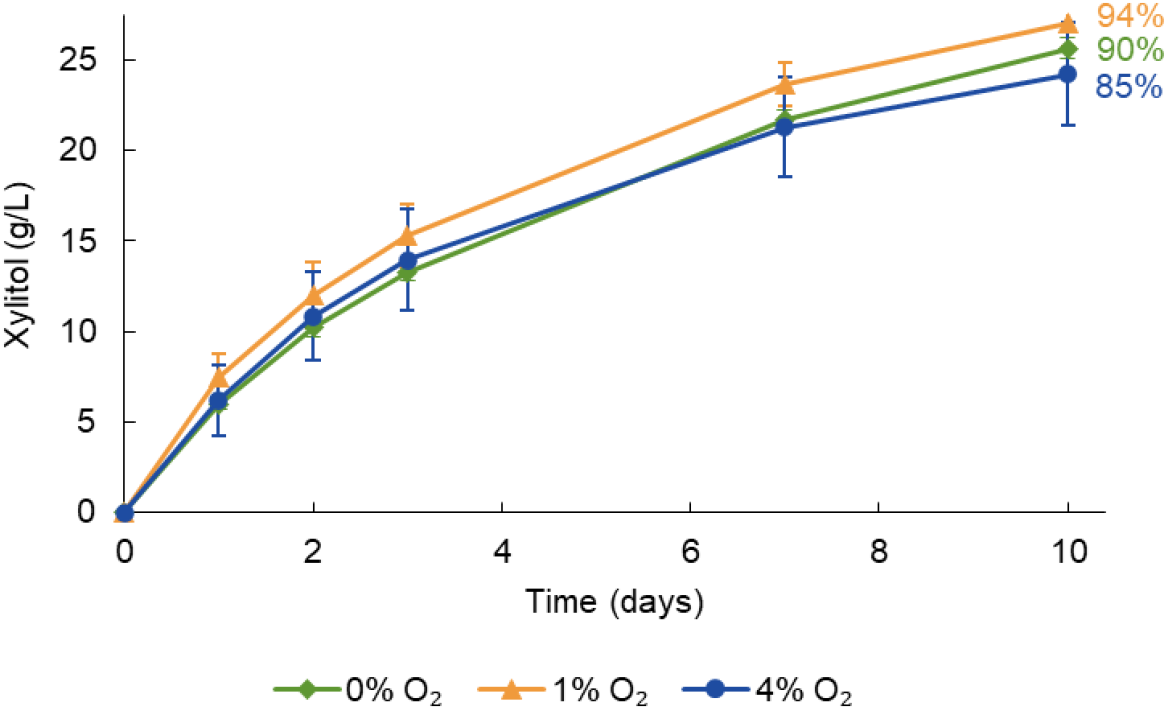
H_2_-driven production of xylitol by *C. necator* △*phaCAB* on different oxygen concentrations. The reaction mixtures contained initially ∼30 g/L of D-xylose in 100 mM sodium phosphate buffer (pH 7.0) with cells at OD_600_ of 16. The used gas mixtures consisted of 0% O_2_ and 100% H_2_, 1% O_2_, 2% N_2_, and 97% H_2_ or 4% O_2_, 13% N_2_, and 83% H_2_. The average from three bioconversions is shown with the standard deviation. The percentages represent the final conversion yields, calculated by dividing the xylitol concentration measured at the final time point by the xylose concentration at the start of the experiment.

### Higher initial sugar concentration can speed up the bioconversion rate

Four different D-xylose concentrations were tested to evaluate their impact on xylitol production rates. With increasing xylose concentration, the rate of conversion increased (Figure 4). The highest xylitol productivity in the first 48 hours (0.7 g L^-1^ h^-1^) was reached with the highest xylose concentration used (114 g/L D-xylose). However, conversion proceeded faster at a lower xylose concentration: 99% of 13 g/L of D-xylose was converted to xylitol in 7 days while 83% of 34 g/L D-xylose was converted within the same time.

**Figure 4.**
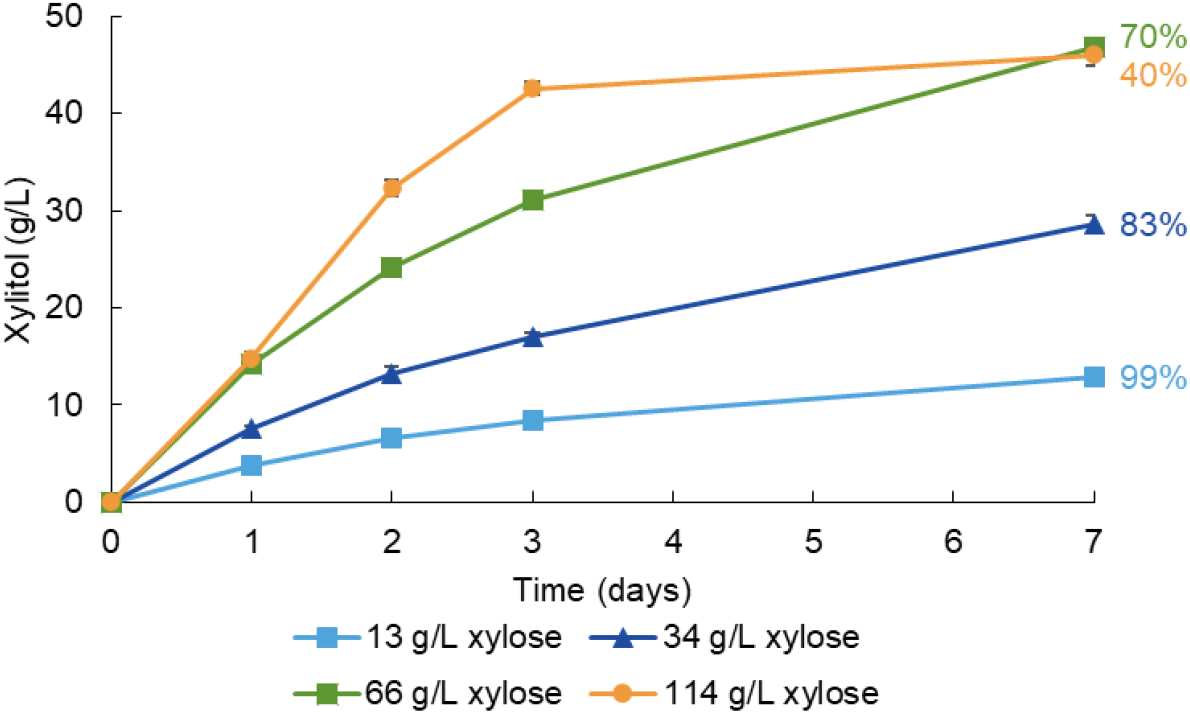
H_2_-driven production of xylitol by *C. necator* △*phaCAB* at different D-xylose concentrations. The reaction mixtures were composed of 13, 34, 66, and 114 g/L of D-xylose in 100 mM sodium phosphate buffer (pH 7.0) and cells at OD_600_ of 13-15. The headspace contained 100% H_2_. The average from three bioconversions is shown with the standard deviation. The percentages represent the final conversion yields, calculated by dividing the xylitol concentration measured at the final time point by the xylose concentration at the start of the experiment.

In samples with the highest xylose concentrations (66 and 114 g/L), the final xylitol concentration reached 46 g/L. We hypothesized that H_2_ in the headspace was limiting and that measuring H_2_ consumption could allow us to estimate the electron conversion efficiency of hydrogen into xylitol. On average, 30 mL of gas was consumed under both conditions, equivalent to 1.2 mmol of H_2_. Given that 0.2 mmol of xylitol was produced without H_2_ (as shown in Figure 2) and 1.3 mmol was produced in total, it can be assumed that 1.1 mmol of xylitol was produced with the help of H_2_ under both conditions. Consequently, more than 90% of the energy derived from H_2_ was spent for the bioconversion, as one mole of H_2_ is required to reduce one mole of xylose (Additional File 1: Full calculations). We wish to highlight that this calculation is an approximation.

### Increased cell concentration enhances the rate of bioconversion

So far, all bioconversions in this study were conducted with a cell concentration range of OD_600_ 13-17. The effect of the amount of the whole-cell biocatalyst was examined at lower and higher cell concentrations: OD_600_ 7 and 60. With the highest cell concentration, 91% conversion of 30 g/L D-xylose to xylitol was achieved in 7 days (Figure 5). Increasing the cell concentration had a positive effect also on the xylitol production rate. In the first 24 hours, the production rates were 0.5 g L^-1^ h^-1^ and 0.1 g L^-1^ h^-1^ for the highest and lowest cell concentrations used, respectively.

**Figure 5.**
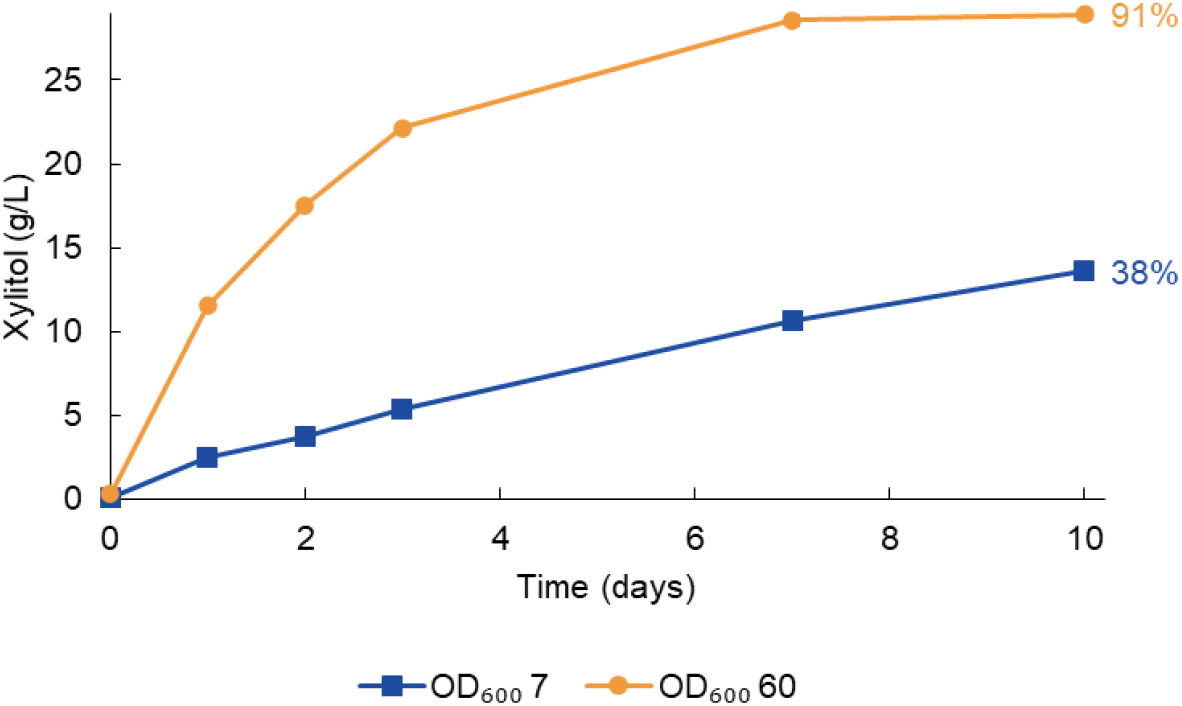
H_2_-driven production of xylitol by *C. necator* △*phaCAB* at different cell concentrations. The initial concentration of D-xylose was ∼30 g/L. The conversions were carried out in a 100 mM sodium phosphate buffer (pH 7.0) under 100% H_2_. The average from three bioconversions is shown with the standard deviation. The percentages represent the final conversion yields, calculated by dividing the xylitol concentration measured at the final time point by the xylose concentration at the start of the experiment.

### Arabinose and ribose are reduced to sugar alcohols by the resting cells

*S. stipitis* XR is also known to convert L-arabinose and D-ribose into their respective sugar alcohols (33). Therefore, we examined whether *C. necator* harboring the xylose reductase could also be used as a biocatalyst for these conversions. Both sugars were successfully reduced to their corresponding sugar alcohols, with the production rates of L-arabitol and ribitol being only slightly lower than those for xylitol (Figure 6).

**Figure 6.**
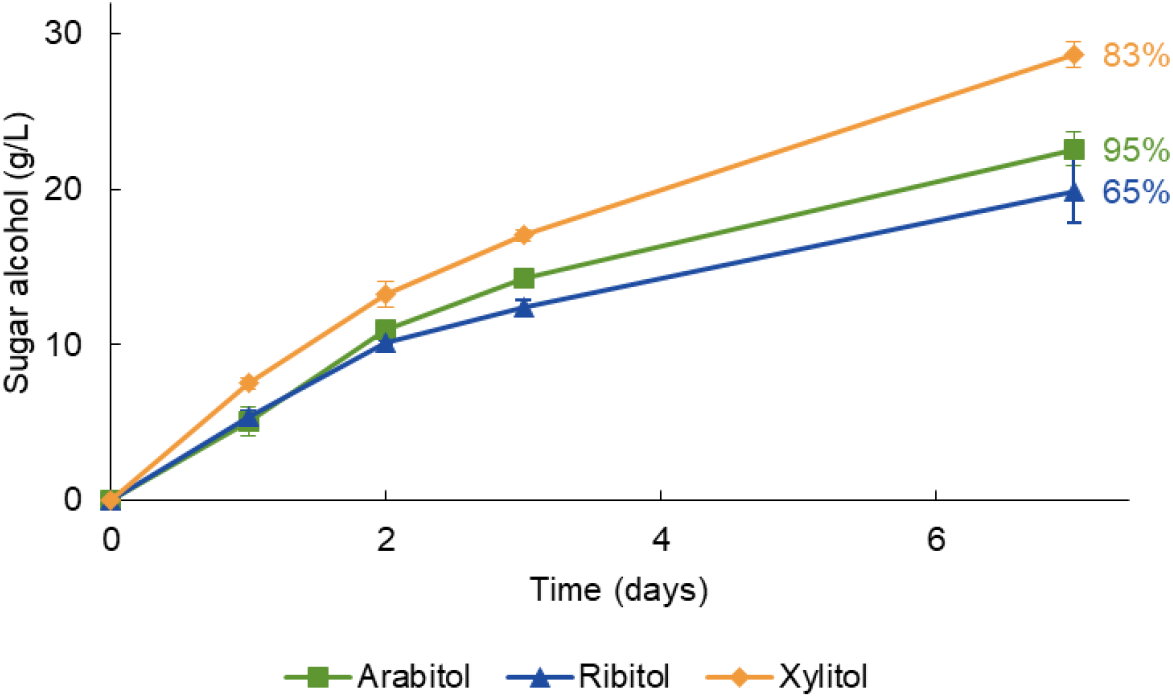
H_2_-driven production of L-arabitol, ribitol and xylitol by C. necator △*phaCAB*. The initial concentrations of L-arabinose, D-ribose and D-xylose were 24, 30 and 34 g/L, respectively, and the corresponding cell densities (OD_600_) were 15, 15, and 13. The experiments were carried out in a 100 mM sodium phosphate buffer (pH 7.0) under 100% H_2_. The average from three bioconversions is shown with the standard deviation. The percentages represent the final conversion yields, calculated by dividing the xylitol concentration measured at the final time point by the xylose concentration at the start of the experiment.

## Discussion

This study aimed to advance the development of *C. necator* as a H_2_-driven whole-cell biocatalyst. We demonstrated nearly complete reduction of 30 g/L of D-xylose to xylitol with 97% conversion yield in resting cells using H_2_ for cofactor regeneration (Figure 2). The effects of different parameters on xylitol production were studied to improve the production rates. Additionally, it was shown that the system can convert L-arabinose and D-ribose into their respective sugar alcohols (Figure 6).

*C. necator* accumulates polyhydroxybutyrate (PHB) as a carbon and energy storage compound. To avoid this accumulation, which could also lead to cofactor oxidation, PHB-negative strains △*phaCAB* and △*phaC* of *C. necator* were used as hosts. Comparison of the △*phaCAB_*xr *and* △*phaC_*xr strains revealed that deletion of the whole PHB pathway (*phaCAB*) enhanced both the bioconversion rate and yield (Figure 2). In the △*phaC* knock-out strain, it is possible that some of the reducing equivalents from H_2_ were consumed by the NAD(P)H-utilizing acetoacetyl-CoA reductase (PhaB) of the PHB pathway. The advantage of knocking out more than just the *phaC* gene has also been observed in earlier studies in non-resting cells of *C. necator*. For instance, complete deletion of the PHB pathway was found to be beneficial for resveratrol production in *C. necator*, whereas deletion of only the *phaC* gene did not improve the titer compared to the wild type (34).

Oxygen is essential for ATP production from H_2_ in *C. necator*. Although the bioconversion reaction itself does not require ATP, it was hypothesized that the cells would require some ATP for cell maintenance. However, the results suggest otherwise (Figure 3). This outcome is advantageous for industrial applications, as oxygen-free production mitigates the risk associated with flammability of H_2_-O_2_ mixtures. The experiments also demonstrated efficient transfer of nearly all hydrogen-derived electrons into the product. When enough H_2_ was present in the headspace, nearly full bioconversion of xylose could be demonstrated. However, liquid solubility of H_2_ is low and hence may still limit the rate of conversion. To test this, bioconversions could be performed under elevated pressure, where H_2_ solubility is increased, but unfortunately it was not possible to test this with the current experimental set-up. Additionally, mass transfer of H_2_ to the liquid phase can be significantly improved by using optimized bioreactors equipped with specialized gas spargers and impellers (35).

The most significant improvements in the bioconversion rate were achieved by increasing the cell concentration and xylose concentration. This is not surprising, as a higher cell concentration provides more catalyst for the conversion to occur, and an increased substrate concentration boosts the reaction rate until enzyme saturation is reached. The most efficient xylitol production systems reported in the literature have reached higher xylitol productivity than the 0.7 g L^-1^ h^-1^ observed in this study. Whole-cell systems have reached up to 12 g L^-1^ h^-1^ (36), while *in vitro* systems have achieved 21 g L^-1^ h^-1^ (23). At a similar cell density to that used in this study, recombinant *E. coli* cells, coexpressing a D-xylose reductase and a glucose dehydrogenase, produced xylitol at 6.4 g L^-1^ h^-1^. In this optimized process, a 100% yield was achieved at an initial D-xylose concentration of 200 g/L using glucose for cofactor recycling (23). Factors that could account for this include lower enzyme activities, slower substrate and product transport into and outside of the cell, and particularly the aforementioned poor H_2_ solubility. Observing the SDS-PAGE gel of Jin et al., it seems that their XR level in the cell was much higher than in this study (Additional file 1: Fig. S3). Although one of the strongest promoters currently known for *C. necator* was used in the present study (30, 37), stronger expression of XR could improve the bioconversion rates. The expression systems for *C. necator* need further development to achieve the expression levels obtained with *E. coli*. On the other hand, the xylose and xylitol transport systems of *E. coli* are likely more efficient than those of *C. necator* because *E. coli* can natively grow on xylose whereas *C. necator* cannot. A BLAST search of the *C. necator* H16 genome using the D-xylose specific transport systems of *E. coli* (XylE and XylFGH) yielded no matches, suggesting that *C. necator* lacks xylose-specific transporters. Xylose is likely transported into the cells by a sugar transporter with side activity for xylose. Hence, heterologous introduction of a xylose transporter could be considered for future studies to further improve bioconversion rates.

The specific activity of XR with NADPH was higher compared to NADH (Table 2). The same result has been observed previously by Verduyn et al. (33). Using NADPH-producing SH, instead of the native NADH-producing SH, with NADPH-dependent oxidoreductases could increase the rate of the bioconversion. The NAD^+^-specific SH from *C. necator* has been engineered to also accept NADP^+^, but its NADP^+^-reducing activity would need to be increased (12). Another option is to use NADH-preferring oxidoreductases or to engineer them to have this preference, ensuring high activity towards NADH.

This article presented the first whole-cell, H_2_-driven biocatalysis study using a PHB-negative *C. necator* strain as the host. Direct comparison of this work to the few prior studies is challenging due to differences in experimental setups, product types, enzyme kinetics, and strains used. Oda et al. (15) reported a productivity of 0.9 g L^-1^ h^-1^ for (*R*)-1,2-propanediol, which is within the same order of magnitude as our findings. Whole-cell cofactor recycling using SH has not only been done in *C. necator*. Lonsdale et al. (38) expressed SH from *C. necator* in *Pseudomonas putida* to perform H_2_-driven bioconversion of n-octane to 1-octanol. The cofactor recycling proved to be effective, resulting in a threefold increase in 1-octanol production in the presence of H_2_. However, the yield and rate of bioconversions they achieved were significantly lower than the ones in this study; Lonsdale et al. reported a maximum productivity of 0.01 g L^-1^ h^-1^, about 100-fold lower than the rates we and Oda et al. achieved.

The limited amount of research in this area offers a wide range of opportunities for improving these organisms to perform H_2_-driven bioconversions towards industrial applications. Bioconversion rates can likely be significantly improved using hosts with improved enzyme activities, elevated pressures, higher cell concentrations and optimized bioreactor designs that enhance hydrogen solubility.

## Conclusions

Cofactor recycling via hydrogenases represents a promising alternative for traditional bioconversion systems because of its atom efficiency, lack of by-products and the prospects of H_2_ becoming a renewable platform chemical of the future. This study demonstrated H_2_-driven bioconversion of D-xylose to xylitol in XR expressing *C. necator* strain. 30 g/L of D-xylose was almost fully converted into xylitol by this system. It was shown that nearly all the energy from H_2_ is harnessed by the bioconversion, demonstrating the potential of the *C. necator* system as an efficient H_2_-driven biocatalyst for sugar alcohol production and potentially other products.

## Data availability

The datasets generated and analyzed during the current study are available from the corresponding author on reasonable request.

## Authors’ contributions

All authors designed research. TJ conducted experiments, data analysis, and wrote the manuscript. All authors revised and approved the manuscript.

## Acknowledgements

The authors thank Enrico Orsi for the *C. necator* △*phaCAB* strain, Guillermo Bordanaba Florit for constructing the *C. necator* H16 △*A0006* strain, Victor de Lorenzo’s lab for the SEVA plasmid, and Ton van Gelder for the help with HPLC analyses. We also thank Solar Foods, especially Juha-Pekka Pitkänen, for the fruitful discussions and the bioeconomy research infrastructures of Aalto University for the support.

---

**Additional file 1: Table S1**. Oligonucleotide primers used in the study. **Table S2**. Synthesized xylose reductase gene used in this study originating from *Scheffersomyces stipitis*. **Table S3**. Upstream and downstream regions of *A0006* used to create *C. necator* H16 △*A0006*. **Figure S1**. The growth of *C. necator* strains △*phaCAB* and △*phaC* on different sugars and sugar alcohols (100 mM). **Figure S2**. Comparison of bioconversion with a single H_2_ flush at the start and H_2_ flush after every sampling. **Figure S3**. SDS-PAGE analysis of soluble extracts by *C. necator* H16 strains. **Full calculations**.

## Additional file

**Table S1.**
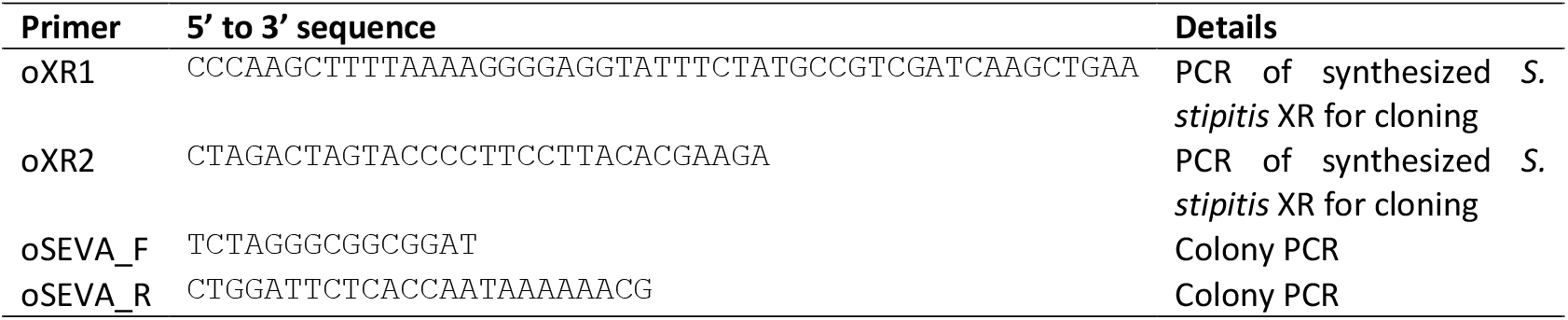
Oligonucleotide primers used in the study.

**Table S2.**
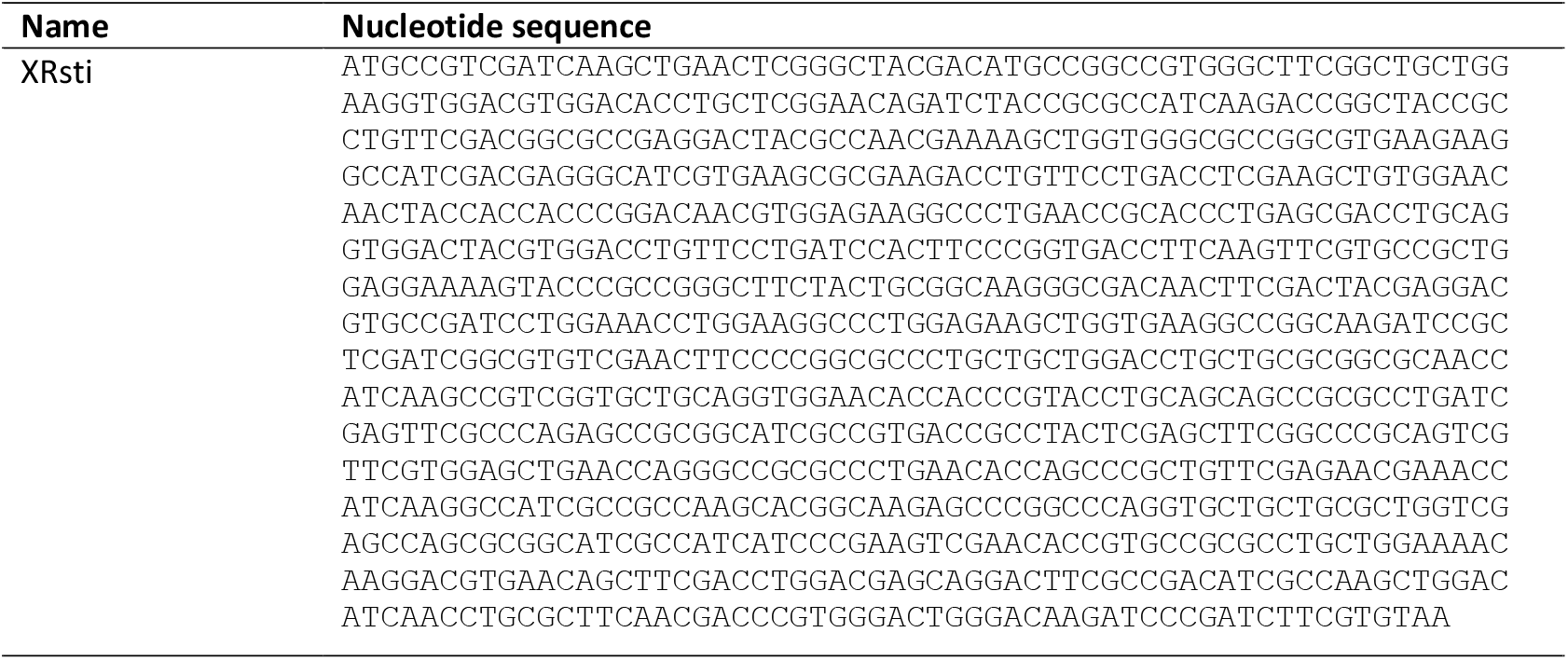
Synthesized xylose reductase gene used in this study originating from *Scheffersomyces stipitis*. Codon-optimized for *C. necator* (957 bp). The original sequence is associated with GeneID 4839234.

**Table S3.**
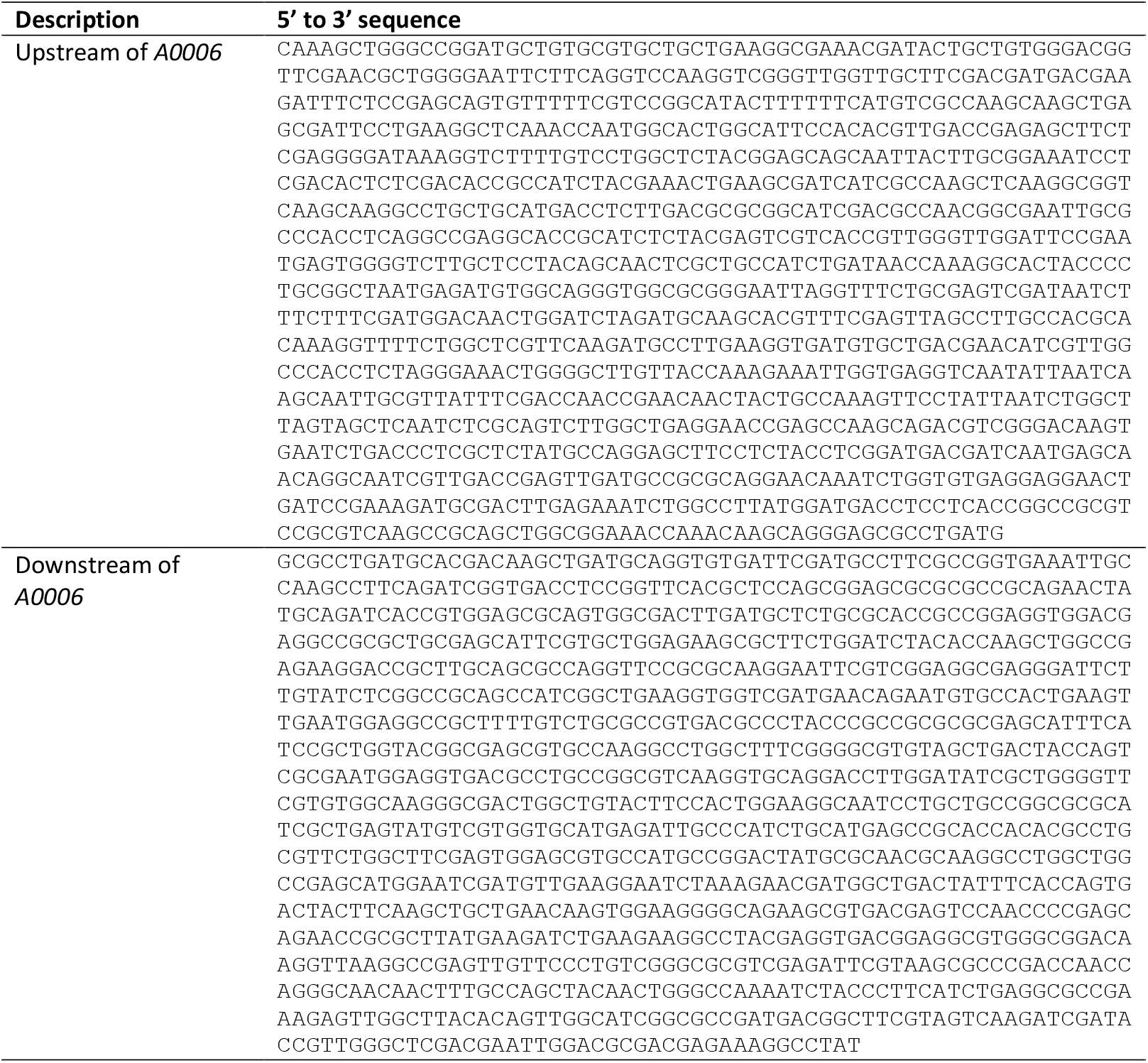
Upstream and downstream regions of *A0006* used to create *C. necator* H16 △*A0006*.

**Figure S1.**
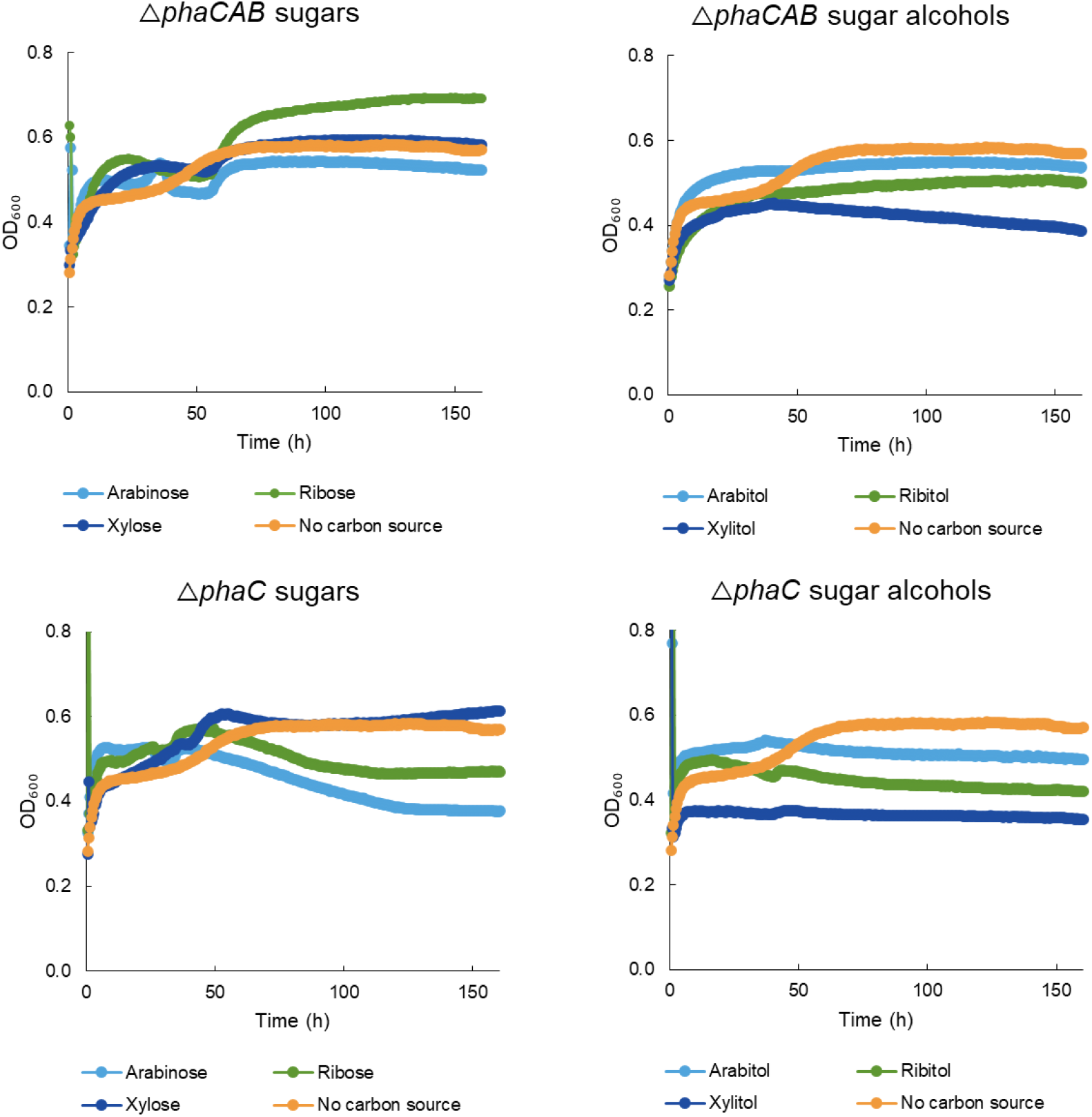
The growth of *C. necator* strains △*phaCAB* and △*phaC* on different sugars and sugar alcohols (100 mM). Precultures were grown in TSB. Cells were cultivated in 96-well plate at an initial OD_600_ of 0.05 in 130 μl of AUT media supplemented with the carbon sources. The cultures were topped with 50 μl of mineral oil to avoid evaporation. The negative control was cultivated in AUT media without any carbon source. The 96-well plate was incubated at 30°C using a double shake orbital at 282 rpm. All conditions had three replicates.

**Figure S2.**
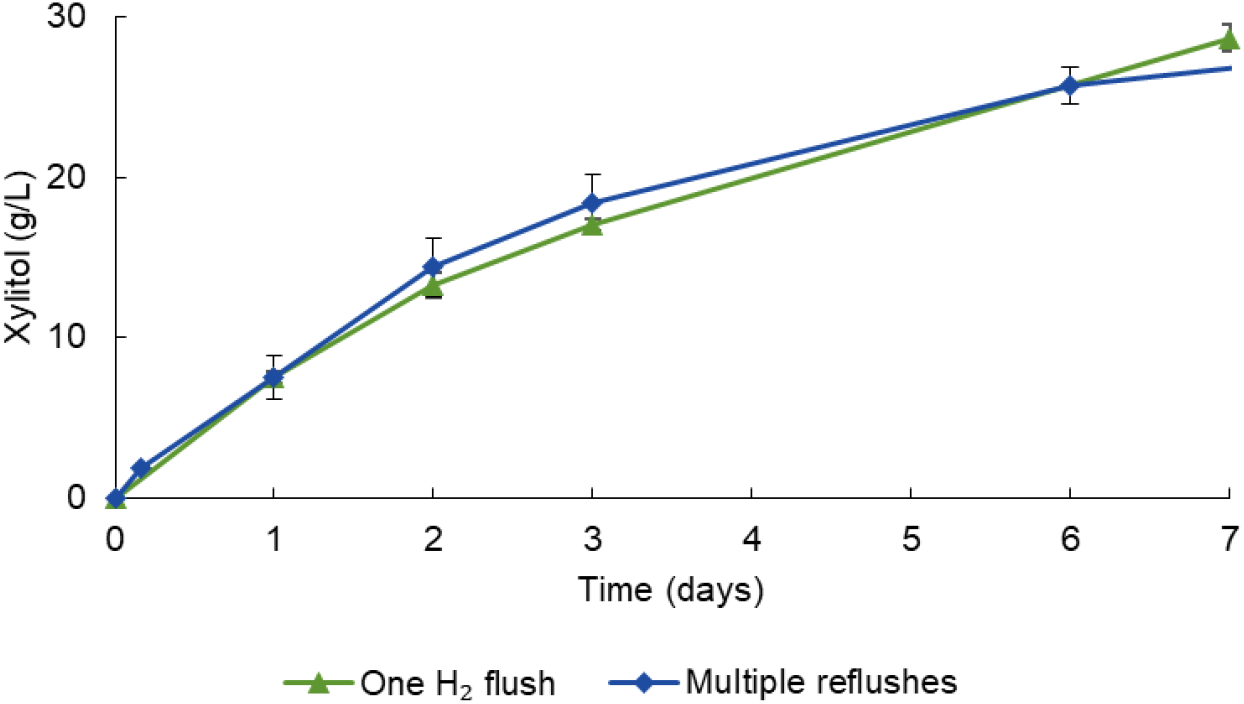
Comparison of bioconversion with a single H_2_ flush at the start and H_2_ flush after every sampling. The initial concentration of D-xylose was 34 g/L (one H_2_ flush) and 30 g/L (multiple reflushes) in 100 mM sodium phosphate buffer (pH 7). The cell concentrations were OD_600_ of 13 (one H_2_ flush) and OD_600_ of 17 (multiple reflushes).

**Figure S3.**
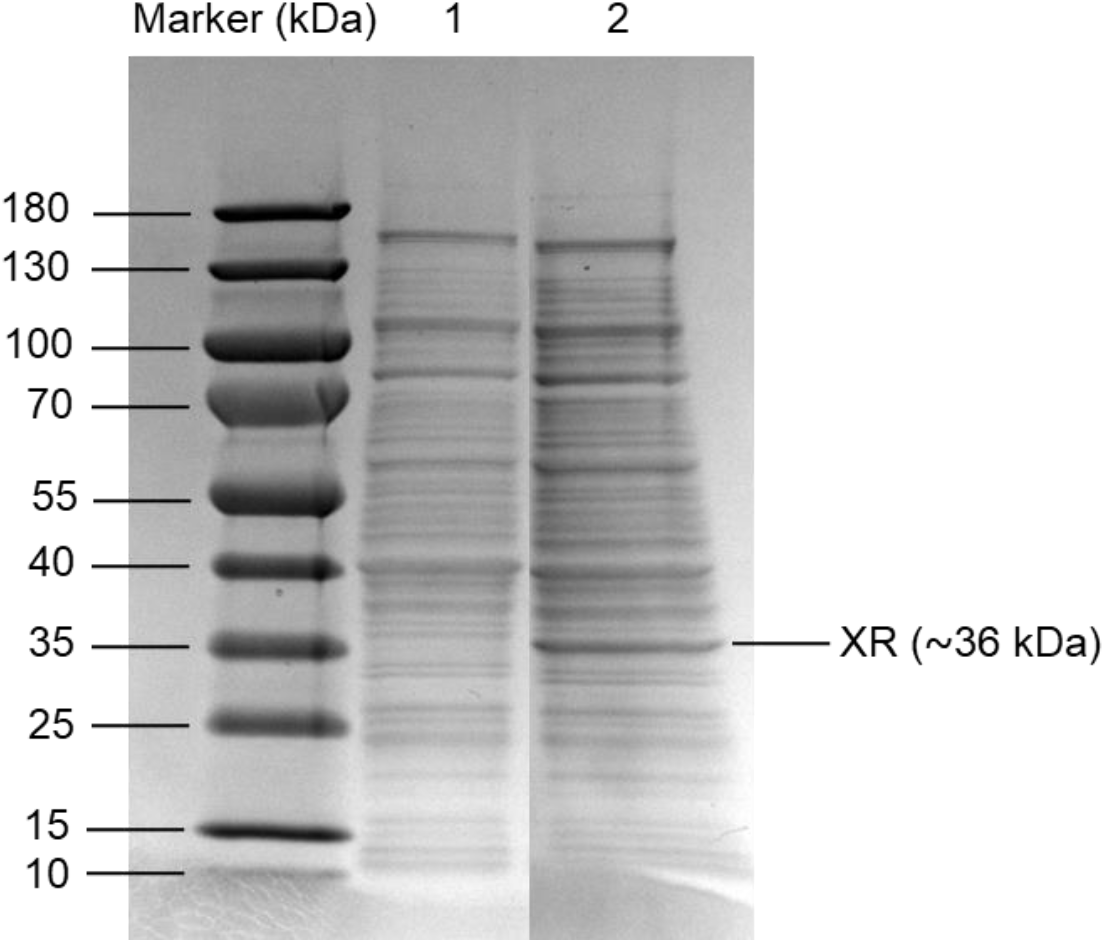
SDS-PAGE analysis of soluble extracts by *C. necator* H16 strains. Lane 1, wild type *C. necator* H16. Lane 2, △*phaCAB*_xr (△*phaCAB* with XRsti under Pj5 promoter).

## Full calculations

### Hydrogen consumption

The ideal gas law was used as a starting point for most of the calculations:

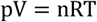

Where p is the pressure (1 bar), V is the gas volume (L), n is the amount of gas (mol), R is the gas constant (0.08314 L bar K^-1^ mol^-1^), and T is the temperature (298.15 K).

The total volume of the serum bottle was 50 mL, with an initial liquid volume of 5 mL. Consequently, the volume of the gas in the bottle was 45 mL, and no overpressure was applied. The initial hydrogen amount in the serum bottles can be calculated:

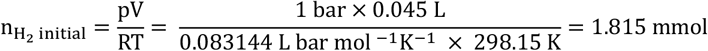

When H_2_ is consumed, only the pressure and the amount of hydrogen in the bottle change, as the gas volume is determined by the fixed volume of the bottle. However, when air is aspirated into the under-pressurized bottle, the volume of aspirated air is equal to the amount of hydrogen consumed. Consequently, the amount of hydrogen consumed can be estimated from the volume of air required to restore the pressure to atmospheric conditions. Additionally, sampling contributed to the under-pressure and must be accounted for:

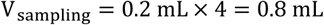

The average aspirated volume into the serum flasks was 30 mL. The consumed H_2_ volume was calculated by subtracting the sampling volume:

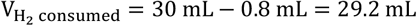

The amount of hydrogen consumed per bottle:

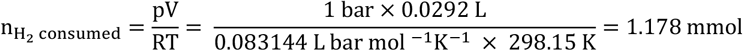

The proportion of the initial H_2_ consumed per bottle is therefore:

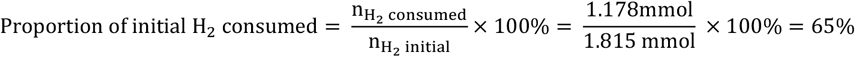

### Xylitol production

The amount of xylitol in the final samples was 46 g/L, which corresponds to 302 mM of xylitol. The final volume in the bottles was around 4.2 mL, considering that 0.8 mL was removed for sampling. Therefore, the final amount of xylitol in the serum bottles was:

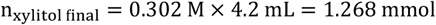

Considering the xylitol in the samples, the total xylitol concentration produced was:

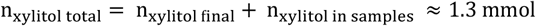

Since some xylitol was produced without the presence of H_2_, this amount must be subtracted from the total to account for xylitol production with H_2_:

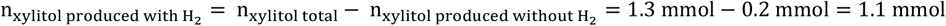

### Amount of H_2_ used for xylitol production from

Since one mole of H_2_ is needed to reduce one mole of xylose, the amount of H_2_ used for xylitol production is 1.1 mmol. The proportion of H_2_ used for xylitol production is:

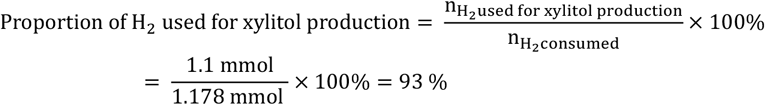

### Error considerations

These calculations are estimates, and several factors may contribute to errors:

- The aspirated volume was measured using a 50 mL syringe, which lacks a fine-scale resolution.
- Sampling was performed with a 1 mL syringe, which also has limited scale resolution.
- Inaccurate sampling volumes can affect the total calculated amount of xylitol produced.
- The xylitol amount in samples is an average from multiple conversions.
- The xylitol amount which was produced without H_2_ is also an average from multiple conversions.

